# Accurate and Fast Protein Acylation Identification by Eliminating Position Effects of Cyclic Immonium Ions with Stepped HCD

**DOI:** 10.1101/2025.07.03.662920

**Authors:** Zhao-Yu Zhu, Peng-Zhi Mao, Ching Tarn, Yong Cao

## Abstract

Mass spectrometry (MS)-based proteomics is indispensable for studying post-translational modifications (PTMs). Cyclic immonium (CycIm) ions serve as invaluable diagnostic markers for lysine acylations, yet the principles governing their generation efficiency are poorly understood. Here, we systematically investigate this question and uncover a robust “position effect”: the generation of immonium ions is strongly favored when the modified residue is located near the N-terminus of a tryptic peptide. Utilizing LysargiNase digestion and isotope-labeled synthetic peptides, we demonstrate that this effect is likely driven by the inherent instability of b-type fragment ions during collision-induced dissociation. Furthermore, we show that a stepped HCD strategy enables enhanced sequence coverage and robust CycIm ion detection (∼99%), thereby improving the depth, reliability and speed of PTM identification. Collectively, this work provides fundamental understanding of immonium ion formation and establishes an optimized acquisition and analysis strategy that enhances the efficiency and confidence of PTM analysis.

## Introduction

Protein lysine acylation represents a diverse and critically important class of post-translational modifications (PTMs) that profoundly expands the functional capacity of the proteome and serves as a key regulatory hub in cellular signaling. By covalently attaching various acyl groups, ranging from simple acetyl moieties to complex long-chain fatty acids, to the ε-amino group of lysine residues, cells dynamically modulate protein charge, conformation, stability, subcellular localization, and interaction networks^1–4^. This regulatory versatility is fundamental to orchestrating a vast spectrum of biological processes, including the epigenetic control of gene expression via histone acetylation^5^, the direct linking of metabolic status to enzyme activity through succinylation and lactylation^6, 7^, the modulation of neuronal functions influenced by crotonylation^8^, and the control of protein membrane association and trafficking via fatty acylation^9^.

Mass spectrometry (MS)-based bottom-up proteomics has emerged as the indispensable powerhouse for the discovery and large-scale identification of protein lysine acylation sites within complex biological systems^6, 10–12^. Its paramount importance stems from the fundamental principle that each acyl modification imparts a specific and predictable mass shift to the lysine residue (e.g., +42.01 Da for acetylation, +100.02 Da for succinylation). This characteristic mass change is readily detectable by high-resolution mass spectrometers following proteolytic digestion of proteins into peptides. Consequently, MS offers a highly versatile and unbiased approach capable of identifying a wide spectrum of both well-established and novel lysine acylations simultaneously, without the a priori requirement for modification-specific affinity reagents like antibodies. This inherent genericity allows for comprehensive acylome profiling, enabling researchers to map modification sites with high precision, quantify their abundance, and uncover the dynamic interplay of diverse lysine acylation events that regulate cellular processes.

Building upon the established power of MS-based bottom-up proteomics for identifying protein lysine acylations via their characteristic mass shifts, a significant challenge remains in achieving high-confidence PTM site localization. This requires unambiguous peptide-spectrum matching (PSM), particularly around the modified residue, which can be demanding. To enhance identification accuracy and streamline analysis, the utilization of modification-specific diagnostic fragment ions within MS/MS spectra has proven highly valuable. These ions serve as direct spectral evidence for the presence of a particular PTM, complementing the information derived from peptide backbone fragmentation^13–15^. For instance, diagnostic oxonium ions (e.g., m/z 138.05, 204.09) derived from glycan moieties are widely exploited to trigger targeted data acquisition or filter candidate spectra in glycoproteomic workflows^16, 17^. Furthermore, specific immonium ions, such as those differentiating mono-/di-methylated lysine (m/z 98.07, 112.09) from acetylated lysine (m/z 126.09), have facilitated the discovery of novel histone PTM sites^18–20^. Similarly, the distinctive cyclic immonium ion of lactyllysine (m/z 156.10) has been instrumental in revealing numerous non-histone substrates within the human proteome^21^. Large-scale studies, such as the ProteomeTools project utilizing synthetic peptides, have systematically cataloged diagnostic ions associated with numerous common PTMs on lysine and arginine under various fragmentation conditions (e.g., HCD, ETD)^22, 23^. The detection of these diagnostic ions significantly increases confidence in PTM assignments and helps mitigate false-positive results. However, a systematic understanding of the factors governing the formation of these diagnostic ions – specifically, whether all modified peptides generate them and under which precise conditions (peptide sequence context, modification position, fragmentation energy) – remains largely unexplored, limiting their universal application.

In this study, we systematically examined the factors influencing immonium type diagnostic ion formation. By segmenting peptides according to the modified residue’s location (N-terminal, central, C-terminal) and assessing detection rates and intensities, we uncovered a significant “position effect”: immonium ion generation is markedly enhanced when the modified residue is near the peptide N-terminus. This appears to be a general principle, independent of amino acid or modification type. Further investigation with LysargiNase digestion and isotope-labeled peptides indicates this effect is likely driven by b-type fragment ion instability under CID. We also demonstrate that a stepped HCD strategy improves both sequence coverage and CycIm ion detection (∼99%), thereby increasing the depth, reliability and speed of PTM identification.

## Methods

### Statistical Analysis of Position Effects of Immonium Ions

The raw data were first analyzed using pFind 3^24^ with the following settings: carbamidomethylation on cysteine was added in fixed modifications; oxidation on methionine, protein N-terminal acetylation, and the acylation modification under study were added in variable modifications; MS1/MS2 error tolerances were set to 10 and 20 ppm respectively; apart from these, all other parameters were kept as pFind defaults. Subsequently, a Python script was employed to statistically analyze the position effects of immonium ions, following these steps: (1) Filtering high-confidence PSMs: Identified peptides were required to have a sequencing ion coverage >0.7, have specific enzymatic cleavage at both N- and C-termini, and contain only one instance of the studied modification or only one amino acid residue generating the immonium ions under study. (2) Grouping by position: Each peptide was divided into three positional groups (left/middle/right) based on whether the studied modification/amino acid in the first, middle, or last third of the peptide sequence. (3) Diagnostic ion detection: Spectra were checked for the occurrence of cyclic immonium (CycIm) or linear immonium (LinIm) diagnostic ions. (4) Relative intensity calculation: The relative intensity of a reporter ion was defined as the ratio of its matched peak intensity to the median intensity of all matched peptide fragment ion peaks.

For the energy optimized dataset, where diagnostic ions triggered MS2 acquisitions at multiple energies, initial identification was also performed using pFind 3. To avoid potential identification bias introduced by differing energy settings, the identified peptide at one energy was propagated across the set of MS2 spectra triggered by the same diagnostic ion at other energies. No sequence coverage filter was applied, allowing analysis of the effects of varying energies on peptide fragmentation beyond their effects on diagnostic ions.

### Synthetic peptides

Crude synthetic peptides were obtained from Anhui Guoping Pharmaceutical Co., Ltd., synthesized with N-terminal Fmoc and internal lysine Boc protection. The peptide was first dissolved in dimethylformamide (DMF) with 2% (v/v) triethylamine (TEA), and the light cross-linker, BS2G-d0, was added at a 5:1 molar ratio to the peptide. The reaction proceeded for 2 hours at room temperature with agitation (1000 rpm). The mono-labeled peptide was desalted via C18 SPE and lyophilized. Next, the N-terminal Fmoc group and Boc protected lysine was removed using a cleavage reagent (95:5 TFA:H₂O, v/v) for 2 hours at room temperature with agitation. The peptide was precipitated twice with pre-chilled diethyl ether and centrifugation (e.g., 15,000 × g, 10 min). The dried peptide was redissolved in DMF with 2% TEA, and the heavy cross-linker, BS2G-d4, was added at the same molar ratio as the first step. This second labeling reaction proceeded for 1 hour at room temperature with agitation. The final product was dried, resuspended in 0.1% (v/v) formic acid, and submitted for mass spectrometry analysis.

### BS2G-d0 and BS2G-d4 Labeled HEK 293T lysate

HEK 293T cell lysate was thawed and clarified by centrifugation (e.g., 15,000 × g, 10 min, 4°C). The soluble proteome was collected and diluted in 50 mM HEPES with 150 mM NaCl (pH 8.0). For labeling, BS2G-d0 or BS2G-d4 was added to separate protein aliquots to a final concentration of 1 mM and incubated for 40 min at room temperature with agitation. Reactions were quenched with 20 mM ammonium bicarbonate for 20 min. Labeled proteins were precipitated with six volumes of pre-chilled acetone at - 20°C for ≥ 2 hours. The pellet was collected by centrifugation, solubilized in 8 M urea, and then alkylated with 10 mM iodoacetamide (IAA) for 15 min in the dark. The urea was diluted to 2 M with digestion buffer, and samples were digested overnight with either trypsin or LysargiNase. Digestions were quenched with 1% (v/v) formic acid. Trypsin-digested peptides were desalted via C18 SPE and lyophilized. LysargiNase-digested peptides were fractionated using a Pierce High-pH Reversed-Phase Kit, and fractions were lyophilized. All dried samples were resuspended in 0.1% (v/v) formic acid for LC-MS/MS analysis.

### The production and enrichment of lactylated peptides

Proteins extracted from HuH-7 cells were reduced with 10 mM TCEP and alkylated with 25 mM CAA (37°C, 30 min). Proteins were then precipitated using four volumes of pre-chilled methanol (−20°C, ≥ 2 h). The air-dried pellet was resuspended in 50 mM NH₄HCO₃ and digested overnight with trypsin (50:1 protein/enzyme, w/w, 37°C). Digestion was quenched with 1% (v/v) TFA. Peptides were desalted using C18 SPE columns (activated: 100% ACN; equilibrated: 0.1% FA; wash: 0.1% FA; elution: 40% ACN in 0.1% FA, v/v) and lyophilized.

Lyophilized peptides were resuspended in IP buffer. Anti-lactylation antibody-conjugated beads (pre-washed with PBS) were added, and the mixture was incubated overnight (4°C, with rotation) to enrich lactylated peptides. Beads were subsequently washed extensively with Wash buffer. Lactylated peptides were eluted three times by incubation with 0.2% TFA (room temperature, 10 min with rotation). Combined eluates were desalted again via C18 SPE as previously described and lyophilized for subsequent analysis.

### LC-MS/MS methods

Liquid chromatography-tandem mass spectrometry (LC-MS/MS) analysis was performed on a Vanquish Neo UHPLC system coupled to an Orbitrap Eclipse Tribrid mass spectrometer (Thermo Fisher Scientific). Peptides were loaded onto a 22 cm × 100 μm C18 column packed with 1.9 μm, 120 Å particles (Dr. Maisch GmbH) and separated at a flow rate of 500 nL/min. The mobile phases consisted of 0.1% formic acid in water (Solvent A) and 0.1% formic acid in 80% acetonitrile (Solvent B). The chromatographic gradient was as follows: 2.1 min at 5% B; a linear increase from 5% to 37% B over 85.9 min; 37% to 50% B over 15 min; and 50% to 99% B over 5.1 min, followed by a re-equilibration period.

For standard data-dependent acquisition (DDA), full MS1 scans were acquired in the Orbitrap from m/z 375-1,500 at a resolution of 120,000. The most abundant precursor ions with charge states 2+ to 6+ were selected for MS2 fragmentation via HCD at a normalized collision energy (NCE) of 30%. Key parameters for MS2 acquisition included an isolation window of 1.6 m/z, an AGC target of 5×10⁴, a maximum injection time of “Auto”, a resolution of 15,000, and a dynamic exclusion duration of 30 s.

For the CycIm ion-triggered acquisition method, a two-stage fragmentation approach was employed. Initially, precursor ions were fragmented via HCD at a fixed NCE of 35%. If the characteristic CycIm ion for glutaryl-lysine (m/z 198.1132) was detected in this initial MS2 spectrum, a second, higher-resolution MS2 scan was triggered. This second scan utilized multiple HCD energies with NCE ranging from 21% to 39%. These triggered MS2 spectra were acquired at a resolution of 30,000 with a first mass set to m/z 110 to enhance detection of low-mass fragment ions.

## Results

### 1. Significant Position Effect Across Six Natural Lysine Acylation

To systematically investigate the factors influencing the formation of CycIm ions, we analyzed the fragmentation patterns of six prevalent and biologically crucial lysine acylations: acetylation^5^, lactylation^7^, succinylation^6^, methacrylation^25^, 2-hydroxyisobutyrylation^26^, and malonylation^27^. The fragmentation scheme of CycIm ions was showed in Figure 1A. Our initial analysis of acetylation data suggested a strong positional dependency, wherein CycIm ion generation is favored for modifications located near the peptide N-terminus.

**Figure 1.**
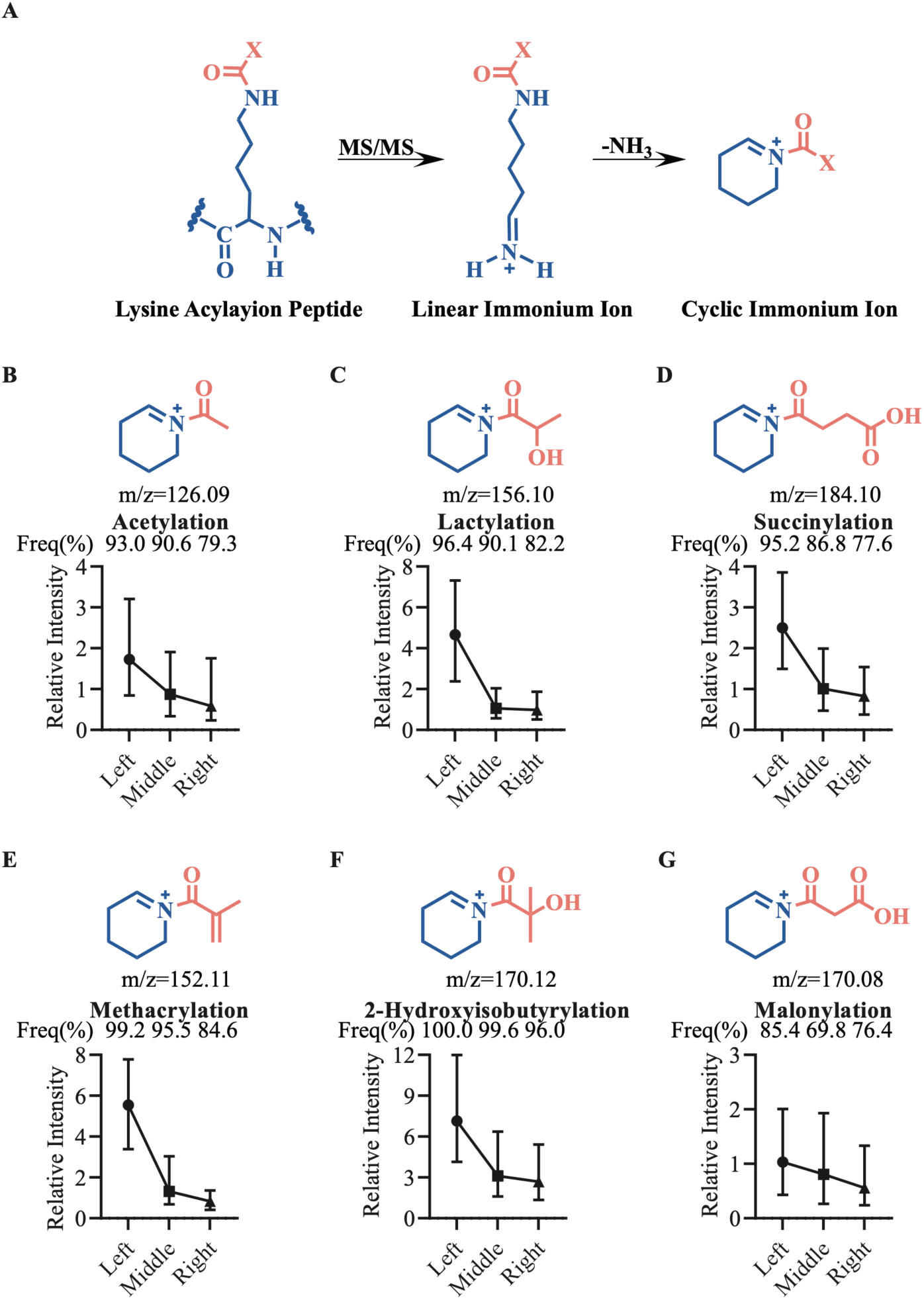
The position effect on CycIm ion generation is general across diverse lysine acylations. (A) Fragmentation scheme illustrating the generation of linear immonium (LinIm) and CycIm ions from an acylated lysine residue. (B-G) Analysis of the detection proportion and relative intensity of CycIm ions for six different lysine modifications. Peptides were categorized based on whether the modified lysine was in the N-terminal, central, or C-terminal third of the sequence. In each panel, the detection proportion for each positional category is shown at the top. The y-axis displays the distribution of CycIm ion relative intensity, with points representing the first quartile (Q1), median (Q2), and third quartile (Q3). The chemical structure for each modification is also shown. Data sources and spectra counts (N) for N-terminal, central, and C-terminal positions are as follows: (B) Acetylation: PXD007630, PXD032953, PXD035832 (N = 776, 1554, 1962) (C) Lactylation: OMIX006151, IPX0003516000, PXD000561, PXD020746, PXD023011 (N = 840, 825, 541) (D) Succinylation: PXD002277 (N = 886, 1685, 1349) (E) Methacrylation: PXD051243 (N = 755, 485, 373) (F) 2-Hydroxyisobutyrylation: PXD038596 (N = 15665, 13506, 4954) (G) Malonylation: PXD035832 (N = 6923, 2431, 608)

To rigorously quantify this observation, we segmented each peptide sequence into N-terminal, central, and C-terminal thirds based on the position of the modified residue. Within each segment, we assessed two key metrics: the proportion of spectra in which the diagnostic CycIm ion was detected and its relative intensity. The results revealed a striking and consistent trend across all six modification types. Both the detection proportion and intensity of CycIm ions progressively diminished as the modification site moved from the N-terminus toward the C-terminus. For instance, with acetylation, the detection proportion decreased from 93% in the N-terminal third to 79% in the C-terminal third; concurrently, the median relative intensity fell threefold from 1.72 to 0.58 (Figure 1B).

This robust positional dependency, which we term the “position effect,” was faithfully replicated for all other modifications analyzed (Figure 1C-G), indicating that it is a fundamental characteristic of the fragmentation process for these acylated residues.

### 2. The Position Effect is a General Principle of Immonium Ion Formation

To confirm the generality of the position effect beyond endogenous modifications, we first examined its presence in datasets of artificially induced lysine acylations. Using N-hydroxysuccinimide (NHS) ester-based chemistry, we generated three distinct acylated peptide populations: propionylated lysine via N-(Propionyloxy) Zsuccinimide, and mono-linked peptides from the cross-linkers DSS and BS2G^28, 29^. Notably, the mono-link from BS2G is equivalent to lysine glutarylation^30^. Consistent with our previous findings, the characteristic CycIm ions from all three artificial modification types exhibited a strong positional dependency, mirroring the trend observed for natural acylations (Figure 2A-C).

**Figure 2.**
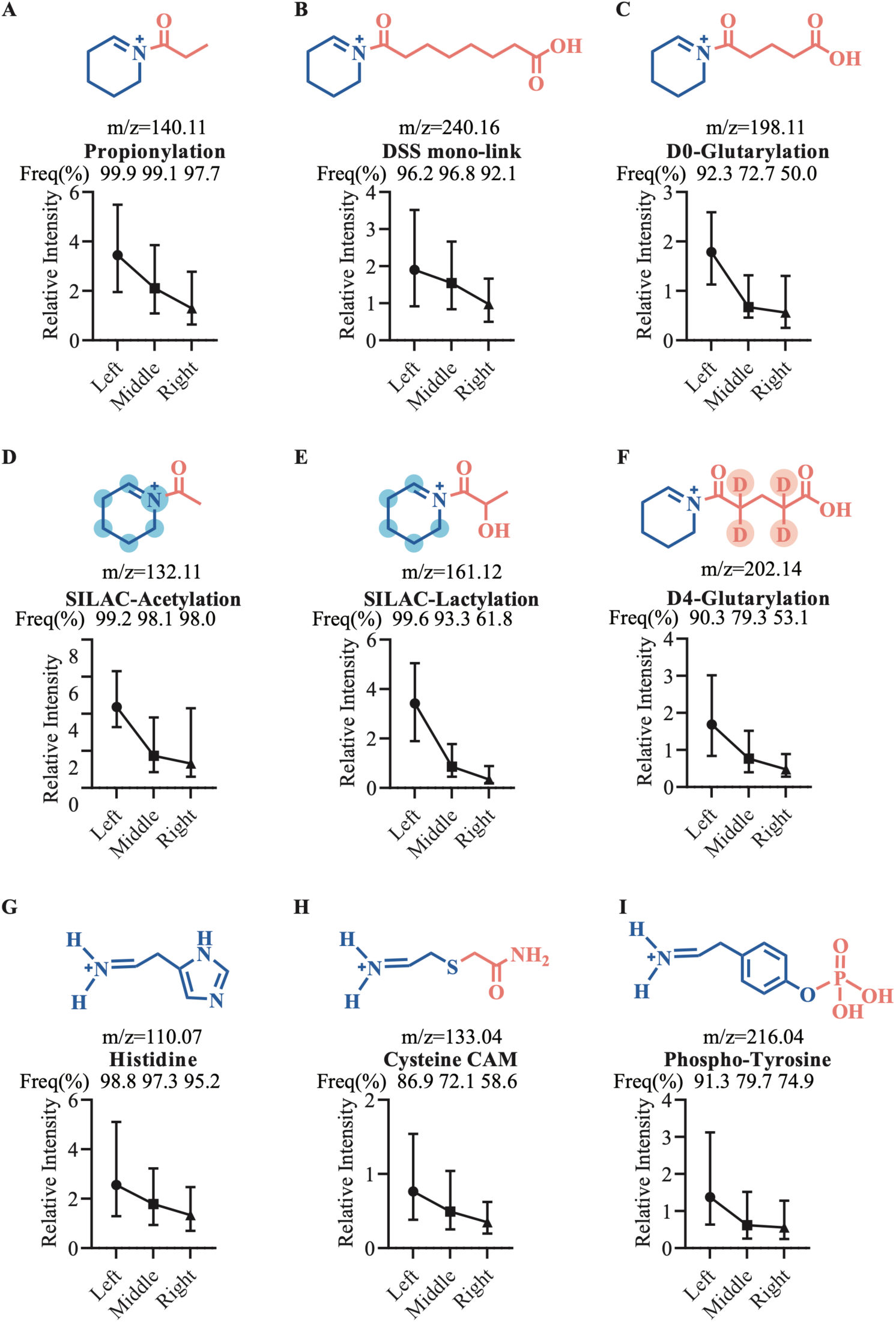
The position effect is a universal phenomenon observed across artificial acylations, isotopically labeled residues, and diverse amino acids. (A-C) Positional analysis of CycIm ions from lysine residues modified with artificial acyl groups: (A) propionylation (via NHS-ester labeling), (B) DSS mono-link, and (C) BS2G mono-link (effectively glutarylation). (D-F) Positional analysis of isotopically labeled CycIm ions, confirming the acyl group as the origin of the additional atoms. Panels show CycIm ions from: (D) acetylation on heavy lysine (¹³C₆¹⁵N₂-Lys), (E) lactylation on heavy lysine (¹³C₆-Lys), and (F) glutarylation via heavy BS2G-d4 labeling. Atoms originating from the isotope labels are highlighted in blue. (G-I) Positional analysis demonstrating the effect extends to LinIm ions from unmodified and other modified amino acids: (G) Histidine, (H) Carboxyamidomethylated Cysteine (CAM-Cys), and (I) Phosphorylated Tyrosine (pY). In each panel, the detection proportion for each positional category (N-terminal, central, C-terminal) is shown at the top. The y-axis displays the distribution of relative ion intensity, with points representing the first quartile (Q1), median (Q2), and third quartile (Q3). Data sources and peptide/spectra counts (N) for N-terminal, central, and C-terminal positions are as follows: (A) Propionylation: IPX0012486000 (N = 2213, 1548, 1652) (B) DSS mono-link: PXD014675, PXD017695 (N = 1275, 1426, 899) (C) BS2G mono-link (Glutarylation): IPX0012486000 (N = 132, 104, 41) (D) SILAC Acetylation: PXD014870, PXD001377 (N = 253, 737, 432) (E) SILAC Lactylation: PXD042051 (N = 234, 236, 89) (F) D4-Glutarylation: IPX0012486000 (N = 139, 96, 51) (G) Histidine: PXD014877, PXD019483 (N = 43190, 38324, 24911) (H) CAM-Cysteine: PXD014877, PXD019483 (N = 114606, 95597, 54162) (I) Phospho-tyrosine: PXD048538 (N = 76006, 90071, 31532)

We then extended this investigation to quantitative proteomics datasets, where stable isotope labels are commonly employed. We analyzed SILAC-labeled (acetylation, lactation) and D4-labeled (glutarylation) datasets to see if the isotopic mass shift would affect this phenomenon. As theoretically predicted, we detected high-frequency CycIm ions with mass increases of +6 Da (or 5 Da) for SILAC and +4 Da for D4. Crucially, these mass-shifted CycIm ions also displayed a clear position effect (Figure 2D-F), demonstrating the robustness of this phenomenon even in quantitative experimental designs.

Finally, to test the hypothesis that the position effect is a fundamental characteristic of peptide fragmentation rather than a feature exclusive to acylated lysine, we broadened our analysis to other residues and modifications. We analyzed the LinIm ions of four unmodified amino acids known for their high fragmentation propensity (histidine, tryptophan, phenylalanine, and tyrosine)^31^. The generation of their LinIM ions, measured by both frequency and median intensity, significantly decreased as the residue’s location shifted towards the C-terminus (Figure 2G, Figure S1A-C). This positional dependency was further confirmed for LinIM ions derived from two common permanent or post-translational modifications: carboxyamidomethylated cysteine and phosphorylated tyrosine (Figure 2H-I). Taken together, these comprehensive analyses strongly support the conclusion that the position effect is a general principle governing the formation of immonium-type fragment ions, independent of the specific amino acid or modification type.

### 3. Direct Validation of the Position Effect Using Isotope-Labeled Synthetic Peptides

To unequivocally demonstrate the position effect while controlling for variables like peptide sequence context and instrument response, we employed a definitive experimental design using synthetic peptides. We synthesized three peptides of varying lengths, each containing two glutarylated lysine (K-Glu) residues at different positions. Crucially, one residue was unlabeled (D0), while the other was heavy-isotope labeled (D4). This strategy enabled the direct comparison of their respective CycIm ions, distinguished by a +4 Da mass shift, within a single MS/MS spectrum (synthesis detailed in Methods; Figure 3A).

**Figure 3.**
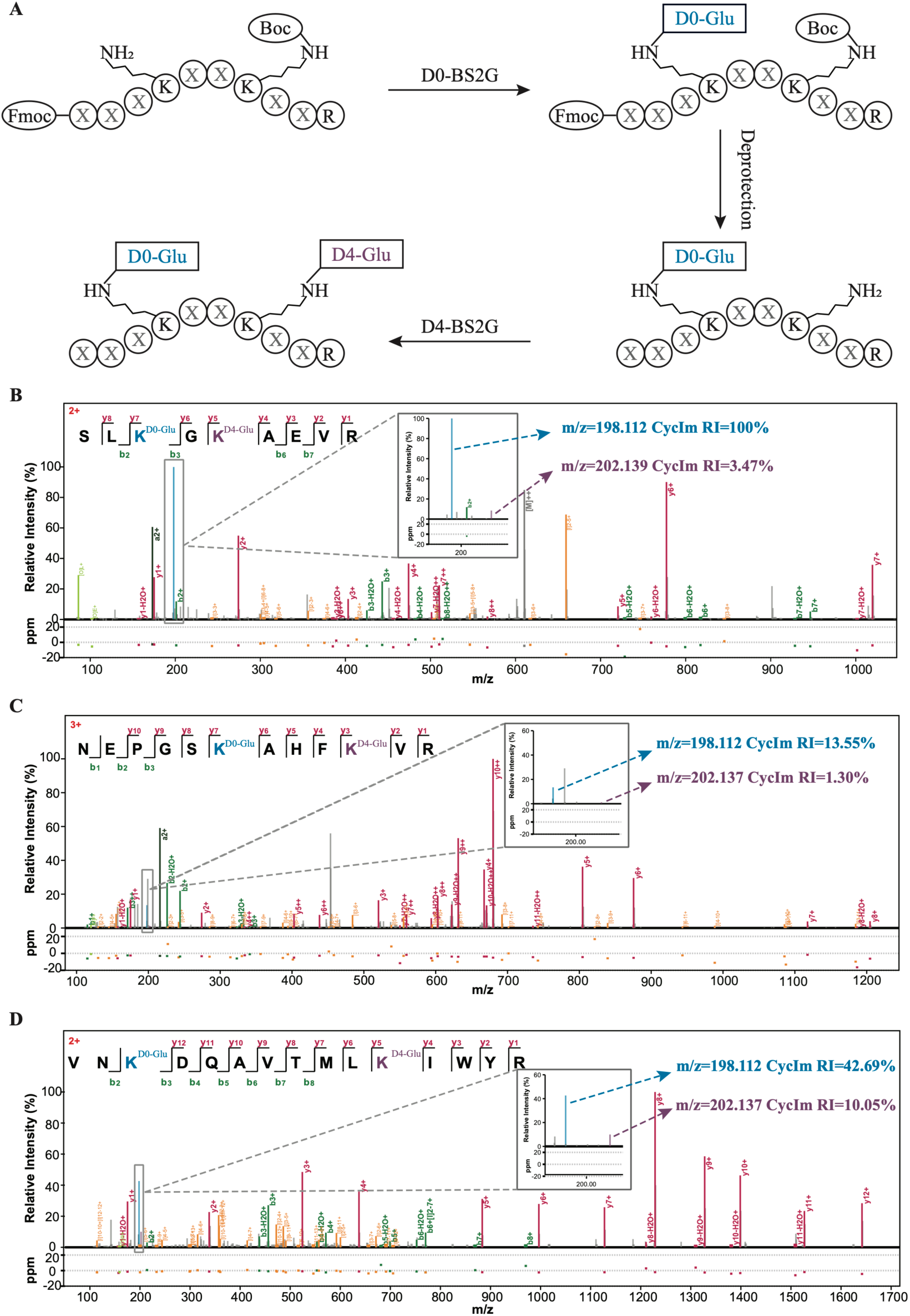
Direct Validation of the Position Effect Using Isotope-Labeled Synthetic Peptides. (A) Schematic of the experimental design for differentially labeling a synthetic peptide with light (BS2G-d0) and heavy (BS2G-d4) cross-linkers. The peptide was synthesized with protecting groups on the N-terminal α-amino group (Fmoc) and the lysine ε-amino group (Boc) to direct the modification. D0-glutarylation and D4-glutarylation are color-coded in blue and purple, respectively. (B-D) Representative MS/MS spectra of peptides labeled with D0-glutarylation and D4-glutarylation. The insets show magnified views of the m/z 200 region, highlighting the CycIm ions for light (blue) and heavy (purple) glutaryl-lysine. The experimentally observed m/z and relative intensities of the D0- and D4-CycIm ions are annotated directly on the spectra.

The results provided unambiguous confirmation of the position effect.

- In a 9-residue peptide **SLK(D0-Glu)GK(D4-Glu)VER**, the K-Glu at the N-terminal position (pos. 3) generated a CycIm ion that was ∼28-fold more intense than the ion from the K-Glu at the central position (pos. 5) (100% vs. 3.5% relative intensity, Figure 3B).
- This trend was maintained in a longer 12-residue peptide **NEPGSK(D0-Glu)AHFK(D4-Glu)VR**, where the central K-Glu (pos. 6) yielded an ion ∼10-fold more intense than the C-terminal K-Glu (pos. 10) (13.5% vs. 1.3%, Figure 3C).
- Similarly, in a 15-residue peptide **VNK(D0-Glu)DQAVTMLK(D4-Glu)IWYR**, the modification closer to the N-terminus (pos. 3) produced a CycIm ion over 4-fold more intense than its C-proximal counterpart (pos. 11) (42.7% vs. 10.1%, Figure 3D).

These experiments, which systematically isolate the positional variable, provide definitive evidence that the proximity of a modification to the peptide’s N-terminus is a primary determinant of its CycIm ion intensity. This validates the position effect as a robust and intrinsic feature of peptide fragmentation, irrespective of peptide length or sequence.

### 4. Reversing the Position Effect by Modulating B-ion Stability

To elucidate the mechanism driving the position effect, we hypothesized that it stems from the fragmentation dynamics of b-ions. B-ions are typically unstable and thus less abundant than y-ions in conventional HCD spectra^32, 33^. A modification near the N-terminus is part of many b-ion precursors (b₂, b₃, b₄, etc.), each of which can fragment to yield a CycIm ion. Conversely, a C-terminal modification is part of fewer b-ions, reducing its chances of being observed.

We designed a critical experiment to test this “b-ion instability” hypothesis: by stabilizing the b-ions, we should be able to alter or even reverse the position effect. To achieve this, we utilized LysargiNase for protein digestion, a method known to enhance b-ion stability by placing a charge-remote basic residue at the peptide N-terminus. As expected, this approach significantly increased b-ion coverage in our glutarylation dataset (Figure S2A-B).

Strikingly, when we re-analyzed the positional dependency in these LysargiNase-generated peptides, the position effect was completely inverted. For all tested immonium ions (CycIm from glutarylation and LinIm from H, W, F, Y, and Cys-CAM), generation efficiency, measured by both detection proportion and intensity, now increased as the residue’s position shifted toward the C-terminus (Figure 4A-C, Figure S2C-E). For glutarylation, the detection proportion surged from 55% (N-terminal) to 92% (C-terminal), with a concomitant threefold increase in median intensity (Figure 4A).

**Figure 4.**
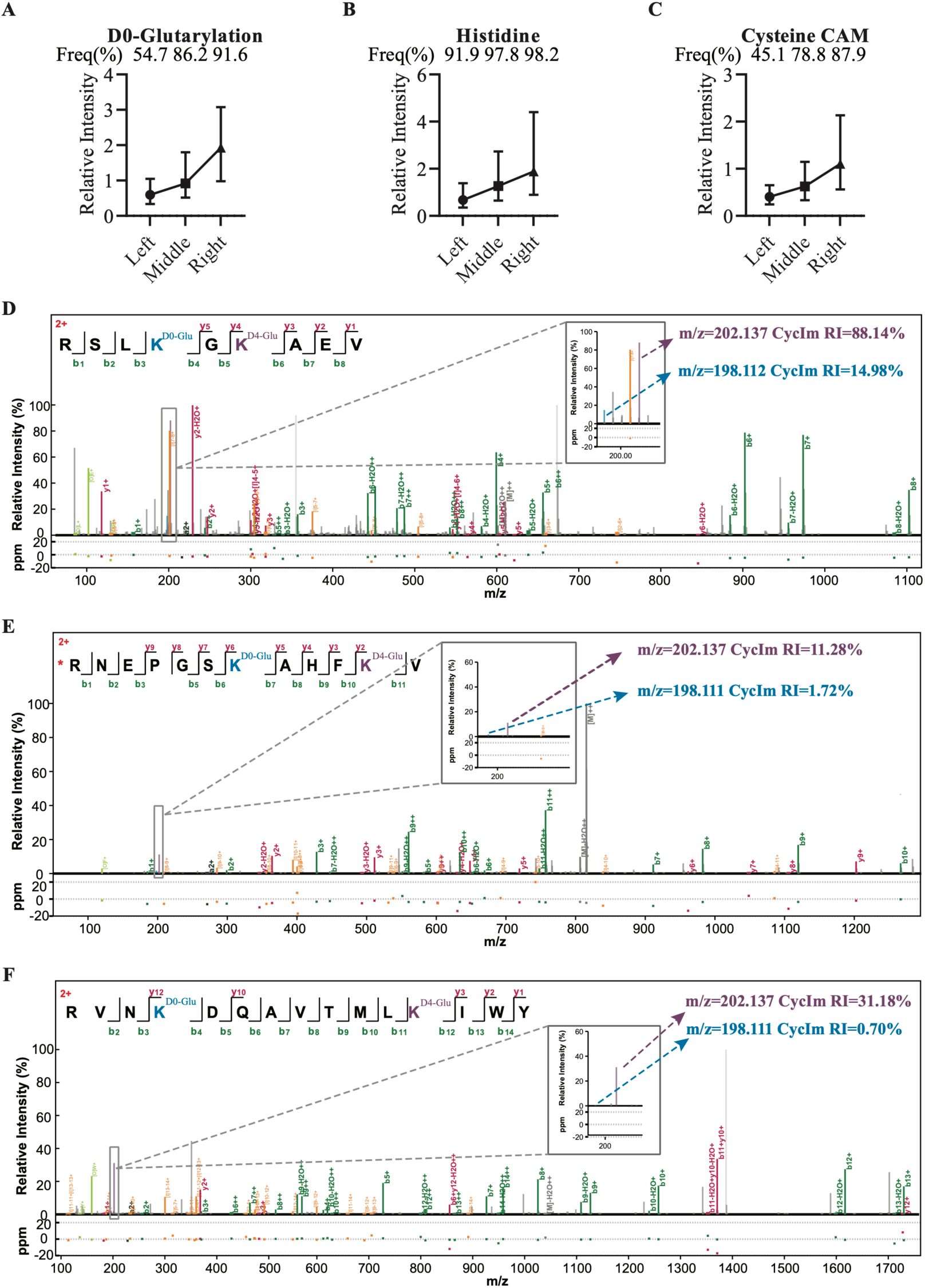
Reversing the Position Effect by Modulating B-ion Stability. (A-C) Positional analysis of immonium-type ions from peptides generated via rAc-Lysarginase digestion. This method produces peptides with a fixed basic residue (Arginine/R or Lysine/K) at the N-terminus, which alters fragmentation behavior. The plot displays the detection proportion (top) and relative intensity distribution (y-axis) for CycIm ions from glutaryl-lysine and LinIm ions from histidine and carboxyamidomethylated cysteine (CAM-Cys) within this specific N-terminally basic peptide population. Data sources and peptide/spectra counts (N) for N-terminal, central, and C-terminal positions are as follows: (A) D0-glutarylation: IPX0012486000 (N = 88, 169, 131) (B) Histidine: IPX0012486000 (N = 701, 905, 1,026) (C) CAM-cysteine: IPX0012486000 (N = 462, 1193, 1325) (D-F) Representative MS/MS spectra of synthetic peptides with an N-terminal basic residue, simultaneously labeled with light (D0) and heavy (D4) glutarylation. The insets show magnified views of the m/z 200 region, highlighting the CycIm ions for light (blue) and heavy (purple) glutaryl-lysine. The experimentally observed m/z and relative intensities of the D0- and D4-CycIm ions are annotated directly on the spectra.

To provide definitive proof of this reversal, we re-synthesized our three model peptides, moving the Arginine residue from the C-to the N-terminus to mimic LysargiNase products. The ratio-metric analysis produced unequivocal results that mirrored the statistical data:

- **RSLK(D0-Glu)GK(D4-Glu)VE** (9-AA): The more C-terminal modification (pos. 7) yielded a ∼6-fold stronger signal than the N-proximal one (pos. 4) (Figure 4D).
- **RNEPGSK(D0-Glu)AHFK(D4-Glu)V** (12-AA): The C-terminal modification (pos. 11) was ∼7-fold more intense than the central one (pos. 7) (Figure 4E).
- **RVNK(D0-Glu)DQAVTMLK(D4-Glu)IWY** (15-AA): The C-proximal modification (pos. 12) produced a signal over 40-fold stronger than the N-proximal one (pos. 4) (Figure 4F).

Collectively, these data strongly support our hypothesis that b-ion stability is the primary mechanistic driver of the position effect. By altering this stability, we can predictably and dramatically invert the phenomenon, thereby providing a clear explanation for this fundamental aspect of peptide fragmentation.

### 5. Stepped HCD Enables High Sequence Coverage and Robust CycIm Ion Detection

Given the diagnostic importance of CycIm ions, we sought to optimize the detection of them. We hypothesized that higher collision energies would favor their formation. However, this immediately presents a potential conflict, as high energies can be detrimental to the sequence-informative b- and y-ions required for peptide identification.

To systematically characterize this trade-off, we implemented a multi-stage fragmentation strategy. Precursors identified as glutarylated (via a CycIm reporter at NCE 35) were re-analyzed across a spectrum of NCEs (range from 21 to 39%). The results confirmed our hypothesis: CycIm ion generation is strongly energy-dependent, with detection rates climbing from 36% to 96% as NCE increased (Figure 5C). In stark contrast, peptide sequence coverage exhibited an optimal range, peaking around NCE 30 before deteriorating at higher energies (Figure 5B). This confirms that a single NCE setting represents a fundamental compromise between identifying the modification and identifying the peptide backbone.

**Figure 5.**
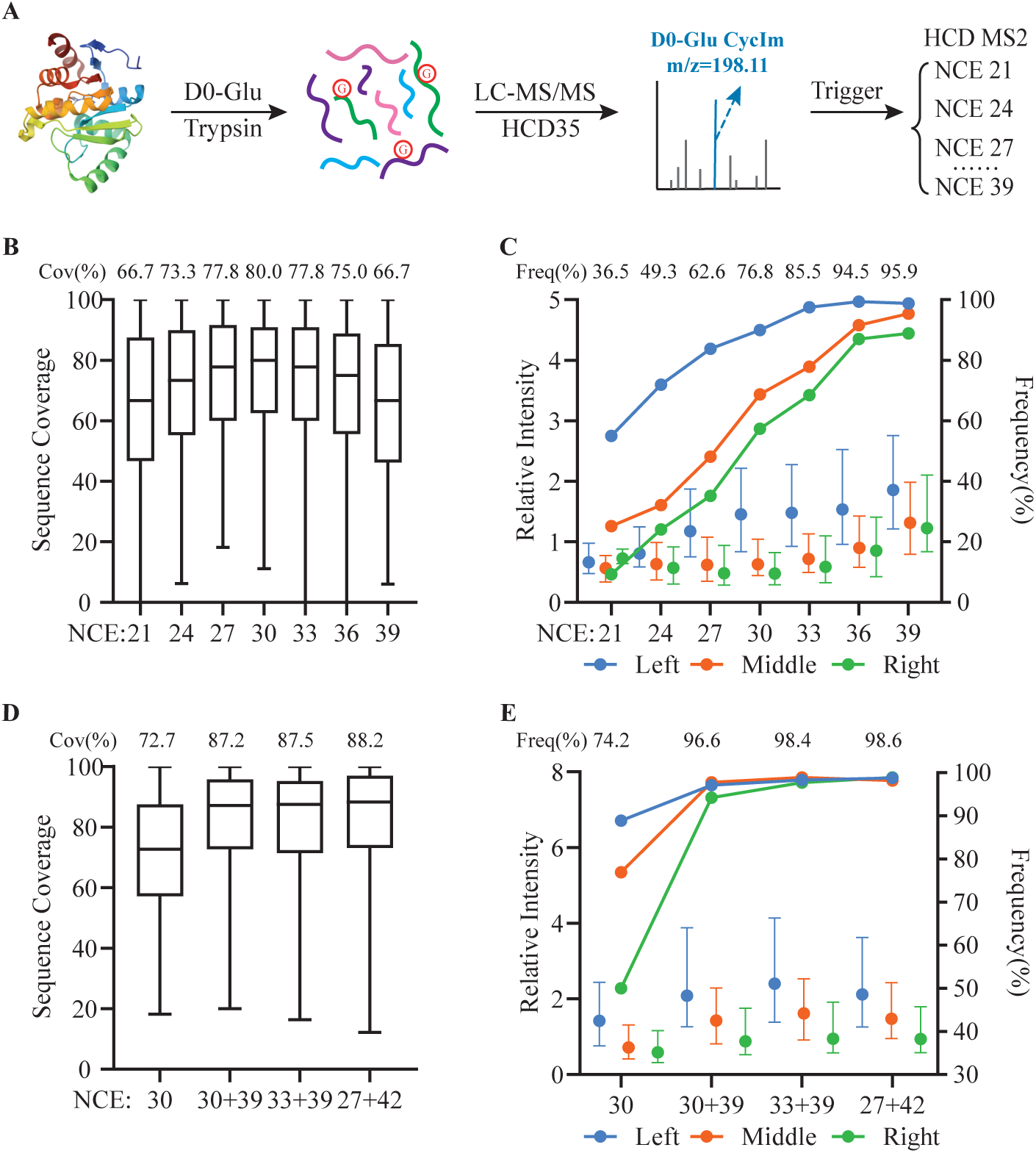
Stepped HCD enhances both peptide sequence coverage and characteristic ion detection for glutarylated peptides. (A) Schematic of a CycIm ion-triggered workflow for optimize collision energy of glutarylated peptides. A fixed collision energy is used (NCE=35), and the presence of the CycIm ion (m/z=198.11) serves as a trigger for multiple energies acquisition. (B, D) Comparison of sequence coverage for glutarylated peptides fragmented with (B) single, fixed normalized collision energies (NCEs) and (D) various stepped HCD strategies. The median sequence coverage for each condition is annotated above the corresponding box plot. The number of spectra (N) was 345 for each fixed NCE level in (B), and 2,355 (NCE 30), 2,247 (NCE 30+39), 2,236 (NCE 33+39), and 2,262 (NCE 27+42) for the conditions in (D). (C, E) Analysis of CycIm ion detection frequency (left y-axis) and relative intensity (right y-axis) for glutarylated peptides fragmented with (C) single, fixed NCEs and (E) stepped HCD strategies. This analysis included only spectra with a sequence coverage greater than 70%. Peptides are categorized by the modification’s position (N-terminal/Left, blue; Central/Middle, orange; C-terminal/Right, green), and the median detection frequency across all positions for each condition is annotated at the top. The number of qualifying spectra (N) for each condition was: (C) The number of spectra (N) was 345 for each fixed NCE level; (E) N = 948 (NCE 30), 1,697 (NCE 30+39), 1,678 (NCE 33+39), and 1,756 (NCE 27+42).

To resolve this conflict, we evaluated stepped HCD as a potential solution to achieve both goals simultaneously. We tested various stepped NCE and compared them to the optimal single-energy settings. The results were compelling: stepped NCE like 30-39 and 27-42 successfully combined the key advantages of both energies. They achieved the high CycIm detection rates and intensities characteristic of a high-energy scan while preserving the high sequence coverage characteristic of a moderate-energy scan (NCE 30) (Figure 5D-E). This optimized fragmentation resulted in obvious improvement in peptide identification, boosting sequence coverage from 72% to 87% and enabling the identification of ∼15% more unique glutarylated peptides (Figure 5B, Figure S3A).

Therefore, we demonstrate that stepped HCD is a superior acquisition strategy for modified peptide analysis, effectively overcoming the inherent trade-off of single-energy fragmentation and enabling confident identification of both the modification and its carrier peptide.

### 6. Apply Stepped HCD on Protein Lactylation Identification

As a recently popularized post-translational modification, protein lactylation provides a direct molecular link between metabolic state, such as glycolytic activity, to the functional regulation of the proteome. This modification has been shown to be a pivotal regulator in a range of fundamental biological processes, including the activation of immune cells, and the mechanisms of DNA repair^34–36^.

To demonstrate the practical utility and robustness of our approach, we applied the stepped HCD strategy to the analysis of endogenous lactylation within a complex cellular proteome. This test yielded significant and multifaceted improvements in both data quality and downstream analytical throughput. Specifically, the method enhanced peptide sequence coverage (89% vs. 80%) while, more critically, increasing the detection rate of the diagnostic CycIm ion to over 99% (Figure 6A-B). This improvement was particularly transformative for peptides with C-terminally located modifications—a group notoriously difficult for diagnostic ion generation—where detection rates surged from a mere 40% to near-quantitative levels (99%) (Figure 6B, 6E-F). This superior fragmentation quality directly translated into a substantial > 50% gain in unique lactylated peptide identifications, enabling a more comprehensive characterization of the lactylome.

**Figure 6.**
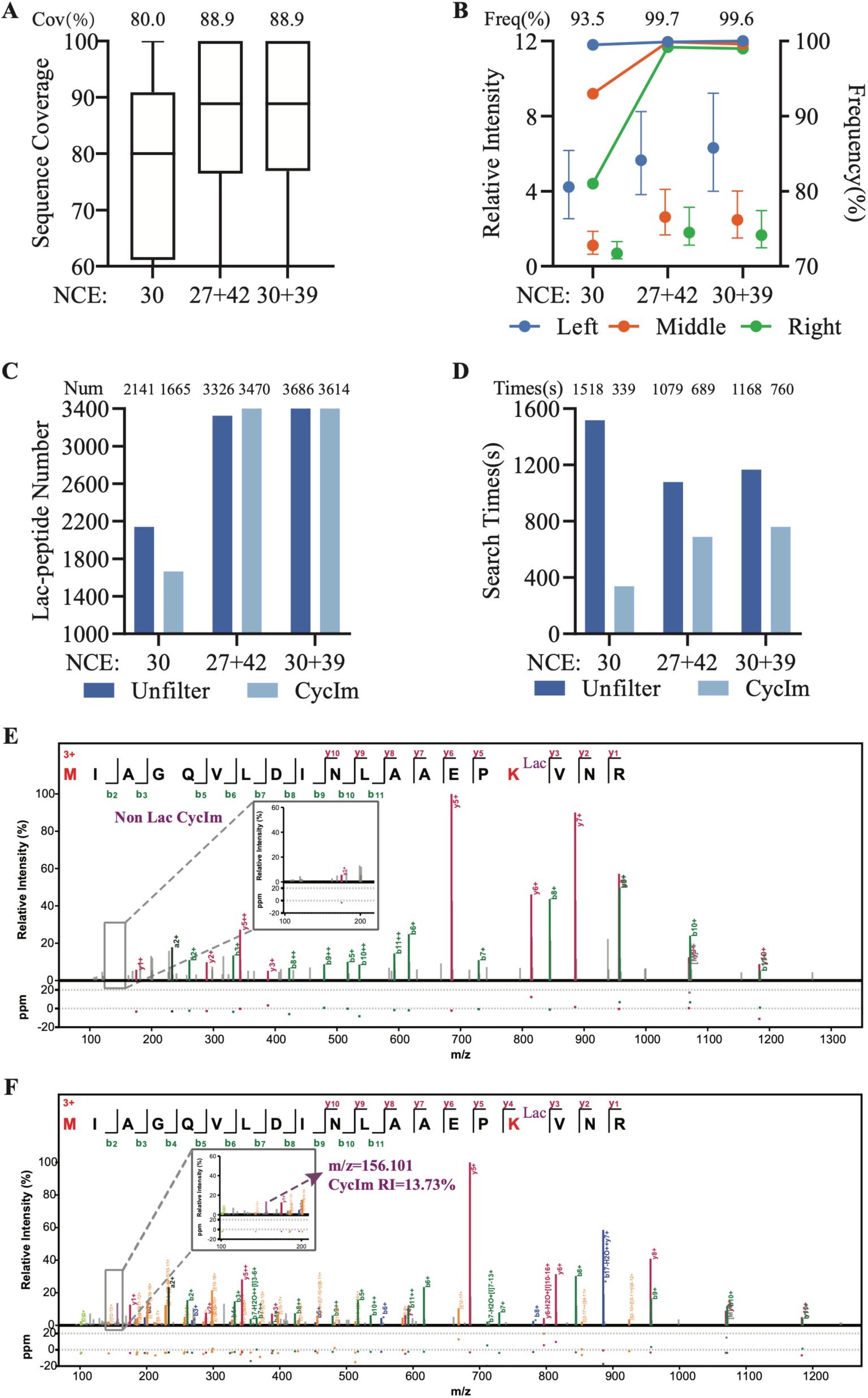
Application of stepped HCD to improve the identification of lactylated peptides. (A) Comparison of sequence coverage for lactylated peptides fragmented with a single fixed NCE (30) versus two different stepped HCD strategies. The median sequence coverage for each condition is annotated above the corresponding box plot. The number of spectra analyzed (N) was 4,551 (NCE 30), 7,069 (NCE 30+39), and 8,254 (NCE 27+42). (B) Analysis of CycIm ion detection frequency (left y-axis) and relative intensity (right y-axis) under the same fragmentation conditions as in (A). This analysis included only spectra with a sequence coverage greater than 70%. Peptides are categorized by modification position (N-terminal/Left, blue; Central/Middle, orange; C-terminal/Right, green), with the median detection frequency across all positions annotated at the top. The number of qualifying spectra (N) was 2,713 (NCE 30), 5,718 (NCE 30+39), and 6,817 (NCE 27+42). (C-D) Evaluation of a CycIm ion-based pre-filtering workflow. Comparison between using the full, unfiltered dataset versus a dataset filtered for spectra containing the lactyl-lysine CycIm ion (m/z 156.10). The plots show (C) the number of identified lactylated peptides and (D) the required database search time. Unfiltered and filtered approaches are color-coded in dark and light blue, respectively. The results of duplicate experiments are combined. (E-F) Representative MS/MS spectra of the same peptide, fragmented with (E) a fixed NCE of 30, where the CycIm ion is absent, and (F) a stepped NCE of 27-42, where the CycIm ion is robustly detected. The insets show magnified views of the m/z 100-200 region, with the lactyl-lysine CycIm ion highlighted in purple.

Furthermore, the high fidelity and near-universal detection of the CycIm ion enabled a powerful pre-filtering strategy. By selectively processing only spectra containing this diagnostic ion, we achieved 35% reduction in database search time while retaining 99.5% of the original identifications (Figure 6C-D, Figure S3A-B). This approach directly addresses the growing computational burden associated with high-throughput proteomics, offering a path to more efficient data analysis with negligible loss of sensitivity.

## Discussion

The discovery and characterization of post-translational modifications (PTMs) by mass spectrometry hinges on interpreting complex fragmentation spectra. While diagnostic ions are invaluable for confident PTM identification, the principles governing their formation have remained largely empirical. A central contribution of this work is the systematic characterization of a potent “position effect,” wherein the N-terminal proximity of an acylated lysine dramatically enhances the generation of its diagnostic cyclic immonium (CycIm) ion.

This principle, while robust, does not operate in a vacuum but within the intricate landscape of gas-phase peptide fragmentation, a process governed by a confluence of factors including precursor charge state, local sequence context, and activation energy. This comprehensive framework explains why the position effect, though a dominant trend, is not absolute. Occasional deviations, such as N-terminally modified peptides yielding suboptimal diagnostic ions, are therefore not contradictions to our central thesis. Rather, they are critical illustrations of the competitive nature of fragmentation, where specific structural features or energetic conditions may favor alternative dissociation channels over CycIm ion formation. Therefore, we propose that the residue’s position introduces a strong, systematic bias that acts as a key predictive factor. This transforms our understanding of diagnostic ion formation from a largely stochastic view to one that incorporates a powerful mechanistic element, enabling a more nuanced and accurate interpretation of quantitative PTM data.

We developed a stepped HCD strategy to address the challenge of acquiring both peptide sequence and modification-specific diagnostic ions within a single fragmentation event. This approach applies combination collision energies during one MS/MS scan, creating a fragmentation environment that is simultaneously optimal for generating extensive b- and y-ion series for sequence determination and for producing robust diagnostic ions (e.g., CycIm) characteristic of a specific modification. The utility of this method has been validated not only for novel PTMs but also for established complex analyses like cleavable cross-linking and glycoproteomics^16,37^. The underlying principle represents a versatile paradigm for any modification analysis where concurrent capture of sequence and diagnostic ions is beneficial, thereby increasing the confidence and depth of PTM studies.

The exceptional data quality afforded by the stepped HCD method—namely, high sequence coverage coupled with near-quantitative diagnostic ion generation (> 99%)— unlocks significant potential for advanced bioinformatic workflows in three aspects: 1) Enhanced computational efficiency: The consistent presence of a diagnostic ion serves as a unique signature that can be used for pre-filtering massive datasets. By rapidly identifying and prioritizing spectra containing this signature, search times can be dramatically reduced, addressing the computational bottleneck created by modern high-speed mass spectrometers. 2) Increased identification confidence: The diagnostic ion provides an orthogonal piece of evidence that can be directly incorporated into peptide scoring models. Integrating diagnostic ion detection and intensity information alongside traditional sequence ion matching scores creates a more robust statistical framework, leading to higher-confidence PTM assignments. 3) Unambiguous site localization: Diagnostic ions can be instrumental in resolving site-level ambiguity. In cases where a modification could plausibly occur on multiple residue types (e.g., an acylation on lysine, serine, or cysteine), the detection of a residue-specific diagnostic ion provides definitive evidence, pinpointing the modification to a specific amino acid type where sequence data alone may be insufficient.

In conclusion, this study bridges a fundamental gap in our understanding of fragmentation by deciphering the ‘position effect’ in immonium ion formation. By translating this insight into an optimized data acquisition method, we not only enhance the analysis of lysine acylations but also pave the way for more sensitive, accurate, and high-throughput characterization of the proteome.

## Supporting information

Supplemental Table 1, and will be used for the link to the file on the preprint site.

## Acknowledgements

We thank Xiao-Hong Xia and Yu-Jiao Qin for their advice in experiment design. The authors gratefully acknowledge financial support from the National Natural Science Foundation of China (NSFC) (No.32401236 to Y. C.).

## Author Contributions

Y.C. conceived the project. Y.C., P.-Z.M., designed the experiments. Z.-Y.Z. executed the wet-lab experiments. P.-Z.M. developed the calculation scripts. Data analysis was performed by Z.-Y.Z., P.-Z.M., and Y.C.. C.T, Z.-Y.Z., P.-Z.M. and Y.C. wrote the manuscript.

## Note

The authors declare no competing financial interest.

## Notes

### Competing Interest Statement

The authors have declared no competing interest.

