## Supplemental Table 1, and will be used for the link to the file on the preprint site. for "Accurate and Fast Protein Acylation Identification by Eliminating Position Effects of Cyclic Immonium Ions with Stepped HCD"

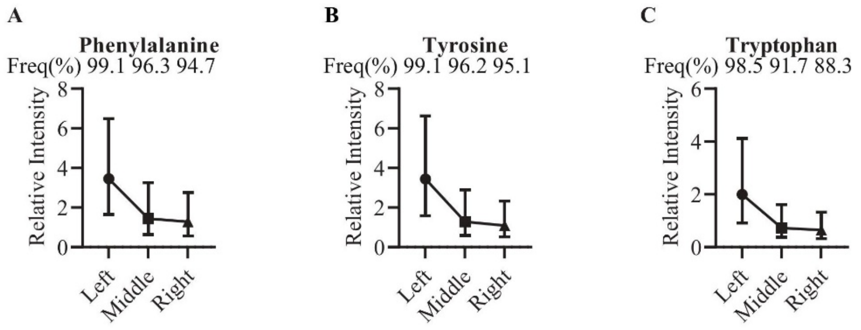


**Supplementary Figure 1. The position effect of LinIm generated from amino acids that are prone to generating LinIm ions**

(A-C) Positional analysis of LinIm ions from peptides generated via Trypsin digestion. This method produces peptides with a fixed basic residue (Arginine/R or Lysine/K) at the C-terminus. The plot illustrates the detection proportion (top) and the relative intensity distribution (y-axis) for LinIm ions of phenylalanine, tyrosine, and tryptophan in peptides that contain only one of these amino acids.

Data sources and peptide/spectra counts (N) for N-terminal, central, and C-terminal positions are as follows:

(A) Phenylalanine: PXD014877, PXD019483 (N = 59,998, 52,054, 32,882)

(B) Tyrosine: PXD014877, PXD019483 (N = 46,107, 40,490, 26,051)

(C) Tryptophan: PXD014877, PXD019483 (N = 19,440, 16,979, 10,861)

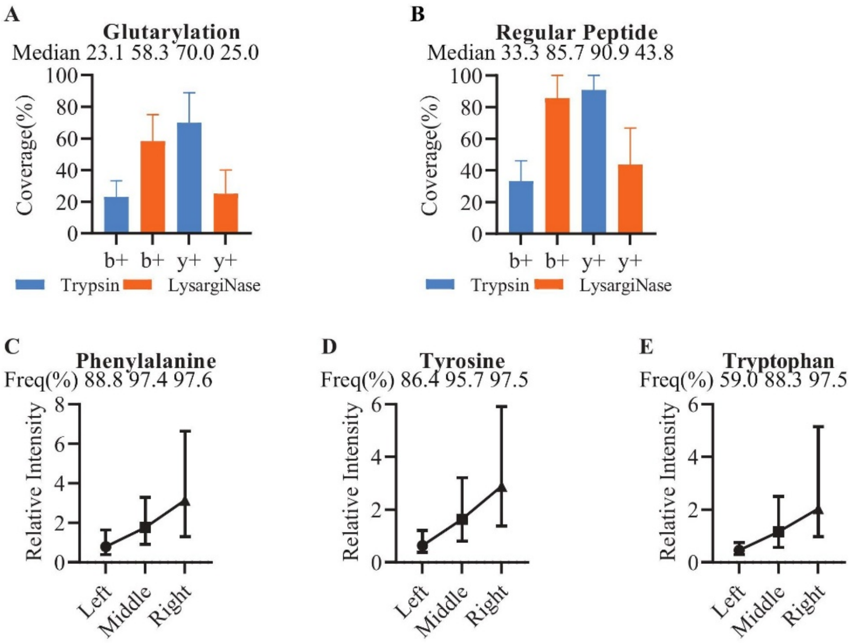


**Supplementary Figure 2. Comparison of the coverage of b ions and y ions and reversed Position Effect of other amino acids via Lysarginase digestion**

(A-B) Compare the sequence coverage of b ions and y ions for glutarylation (A) and other regular peptides(B) under Trypsin or Lysarginase digestion conditions. The plot illustrates the median coverage(top) and sequence coverage (y-axis) of b ions and y ions. All Trypsin digestion peptides have been filtered: the C-terminal is either a lysine (K) or arginine (R) amino acid, while the N-terminal is neither a lysine (K) nor an arginine (R) amino acid. In contrast, the Lysarginase digestion peptides have an N-terminal that is either a lysine (K) or arginine (R) amino acid, and a C-terminal that is neither a lysine (K) nor an arginine (R) amino acid. (C-D) Positional analysis of LinIm ions from peptides generated via Lysarginase digestion. The plot illustrates the detection proportion (top) and the relative intensity distribution (y-axis) for LinIm ions of phenylalanine, tyrosine, and tryptophan in peptides that contain only one of these amino acids.

Data sources and peptide/spectra counts (N) for N-terminal, central, and C-terminal positions are as follows:

(A) Glutarylation: In-house labeled sample (N = 519, 925, 519, 925)

(B) Regular peptides: PXD014877, PXD019483, PXD014574 (N = 102,907, 13,508, 102,907, 13,508)

(C) Phenylalanine: In-house labeled sample (N = 985, 1,195, 1,137)

(D) Tyrosine: In-house labeled sample (N = 803, 952, 901)

(E) Tryptophan: In-house labeled sample (N = 161, 250, 231)

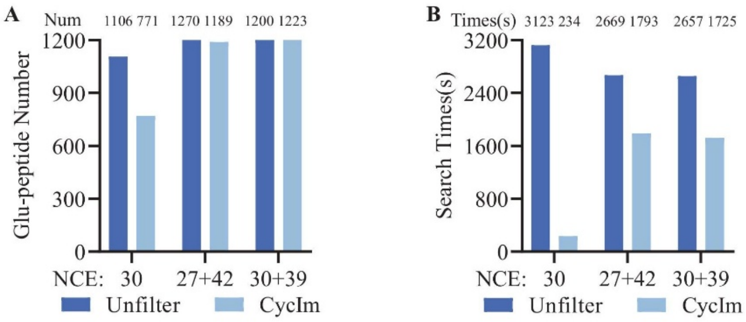


**Supplementary Figure 3. The number of glutarylated peptides and the search times are compared between unfiltered and filtered CycIm ion spectra.**

(A-B) The number and search times of glutarylated peptides are compared between unfiltered and filtered CycIm ion spectra. The plot illustrates the total number and search times in the top. Set two technical replicates for each condition.

Table 1

| Num. | Name | Data Source | Species |
|---|---|---|---|
| Fig.1B | Acetylation | PXD007630[1], PXD032953, PXD035832[2] | Arabidopsis thaliana, Mus musculus, Homo sapiens |
| Fig.1C | Lactylation | OMIX006151[3], IPX0003516000[4], PXD000561[5], PXD020746, PXD023011 | Homo sapiens, Trypanosoma brucei |
| Fig.1D | Succinylation | PXD002277[6] | Homo sapiens |
| Fig.1E | Methacrylation | PXD051243 | Homo sapiens |
| Fig.1F | 2-Hydroxyisobutrylation | PXD038596[7] | Rattus norvegicus |
| Fig.1G | Malonylation | PXD035832[2] | Mus musculus, Homo sapiens |
| Fig.2A | Propionylation | IPX0012486000 | Saccharomyces cerevisiae |
| Fig.2B | DSS | PXD014675[8], PXD017695 | Homo sapiens, Mycoplasma pneumoniae |
| Fig.2C | Glutarylation | IPX0012486000 | Homo sapiens |
| Fig.2D | SILAC-Acetylation | PXD014870[9], PXD001377[10] | Homo sapiens |
| Fig.2E | SILAC-Lactylation | PXD042051 | Homo sapiens |
| Fig.2F | D4-Glutarylation | IPX0012486000 | Homo sapiens |
| Fig.2GH, Fig.S1ABC | Histidine, CAM-Cysteine Phenylalanine, Tyrosine, Tryptophan | PXD014877[11], PXD019483[11] | Homo sapiens, Arabidopsis thaliana, Rattus norvegicus |
| Fig.2I | Phospho-tyrosine | PXD048538[12] | Homo sapiens |
| Fig.3B-D, Fig.4D-F | Prptide | IPX0012486000 | Homo sapiens |
| Fig.4A-C | Glutarylation, Histidine, CAM-Cysteine | IPX0012486000 | Homo sapiens |
| Fig.5B-E | Glutarylation | IPX0012486000 | Homo sapiens |
| Fig.6A-F | Lactylation | IPX0012486000 | Homo sapiens |
| Fig.S2B | Lysarginase | PXD014574 | Homo sapiens |
